# Food preference is associated with distinct large-scale cortical functional connectivity patterns during food-image observation

**DOI:** 10.64898/2026.04.22.720279

**Authors:** Hisato Sugata, Sarang Kim, Takashi Ikeda, Masayuki Hara

**Affiliations:** Graduate School of Welfare and Health Science, Oita University, 700, Dannoharu, Oita, 870-1192, Japan; Faculty of Welfare and Health Science, Oita University, 700, Dannoharu, Oita, 870-1192, Japan; Research Center for Child Mental Development, Kanazawa University, Kanazawa, Japan; Graduate School of Science and Engineering, Saitama University, Saitama, Japan

**Keywords:** food preference, functional connectivity, food cues, alpha band, beta band

## Abstract

Food preference influences behavior toward food-related stimuli, yet the large-scale neural mechanisms underlying this process remain unclear. This study investigated whether preferred and nonpreferred food cues are associated with distinct patterns of cortical functional connectivity during the observation of food images. Data from 25 of the 40 recruited healthy adults were included in the final analysis after excluding individuals with highly unbalanced response tendencies. Participants viewed 150 food images and rated each image on a four-point preference scale. Trials were classified as favorite food (FF) or disliked food (DF). High-density electroencephalography (EEG) was recorded during the task, and source-level ROI-to-ROI functional connectivity was analyzed using amplitude envelope correlation in the alpha (8–13 Hz) and beta (13–25 Hz) frequency bands over the 1000-ms period after food-picture onset. Response time did not differ significantly between FF and DF trials. However, distinct functional connectivity patterns were observed between conditions in both frequency bands. In the alpha band, FF trials involved a network including the cuneus, parietal regions, cingulate regions, and lateral occipital cortex, whereas DF trials involved the isthmus cingulate, caudal middle frontal gyrus, inferior temporal cortex, superior parietal lobule, and lateral occipital cortex. In the beta band, FF trials involved the isthmus cingulate, precuneus, parietal regions, and pericalcarine cortex, whereas DF trials additionally involved frontal regions, including the superior frontal gyrus and pars triangularis. These findings indicate that food preference is associated with distinct large-scale cortical functional connectivity patterns during food image observation, suggesting differential neural processing of preferred and nonpreferred food cues.

## 1. Introduction

Food choice is shaped by multiple interacting factors, including food’s sensory properties, subjective preferences, and the physiological and psychological state of the individual [1, 2]. Clarifying how the brain processes preferred and nonpreferred food cues may therefore provide important insights into the neural mechanisms underlying food selection and motivation to eat [2, 3].

Visual food cues are particularly relevant, as they can influence eating-related decisions before actual consumption [4]. Even in the absence of direct oral stimulation, food images can evoke attentional, motivational, and evaluative processes that contribute to judgments about whether a food is desirable [3, 5]. Previous electrophysiological studies have shown that food-related stimuli modulate oscillatory brain activity, suggesting that the subjective evaluation of food cues is associated with neural dynamics [6].

In particular, low-frequency oscillations provide a useful framework for examining these processes [7, 8]. In particular, alpha-band activity has been linked to sensory processing, perceptual gating, and attentional allocation [7, 9, 10], whereas beta-band activity has often been associated with sensorimotor and evaluative processing [11-13]. Accordingly, oscillatory dynamics in those bands may provide important insights into how food cues are prioritized and evaluated during the early stages of food-related decision-making. A previous EEG study focusing on localized activity changes showed that preferred food images can modulate oscillatory brain activity, supporting the idea that food preference is reflected in the cortical processing of food cues [6]. However, since food preference likely emerges from interactions among visual, attentional, and evaluative systems, a network-level approach may provide more comprehensive information on the neural mechanisms underlying food preference.

The present study investigated the effects of food preference on cortical functional connectivity during food image observation in healthy adults using high-density EEG. By applying source-level functional connectivity analysis, we aimed to characterize the large-scale network organization associated with preferred and nonpreferred food cues. We hypothesized that preferred and nonpreferred food images would be associated with distinct patterns of functional connectivity in the alpha and beta bands, reflecting differences in visual-attentional and evaluative processing related to food preference.

## 2. Methods

### 2.1 Participants

Forty healthy participants were initially recruited. During the task, participants were asked to evaluate food images, indicating whether they wanted to eat the presented item or not. To minimize response bias, participants who selected either “want to eat” or “do not want to eat” in >70% of trials (105 trials) were excluded from subsequent analysis because their responses were considered overly unbalanced. As a result, data from 25 participants were included in the final analysis.

### 2.2 Experimental procedure

Participants were instructed not to eat any food for 3 h before the experiment. Visual stimuli were selected from a standardized food-picture database developed for previous research [14, 15]. A total of 150 food images were presented, consisting of 75 high-calorie (HC) food images and 75 low-calorie (LC) food images. HC images included both sweet and savory foods, whereas LC images included vegetables, fruits, and salads. The selected images were controlled for basic visual characteristics, including color, brightness, spatial frequency, contrast, and visual complexity, and matched in resolution, color depth, background color, and camera distance. All images were presented on a monitor at a viewing distance of ∼50 cm.

The experimental paradigm is illustrated in Figure 1. Each trial consisted of three phases, as in our previous study [6]. First, during the rest phase, a black fixation cross was presented at the center of the screen for 3,000 ms. Thereafter, in the observation phase, a food image was presented for 3,000 ms during which participants were instructed to continuously observe the image. Finally, in the response phase, participants rated their subjective preference for the presented food using a 4-point scale, ranging from 1 (“don’t want to eat”) to 4 (“want to eat”), by pressing one of four buttons.

**Figure 1.**
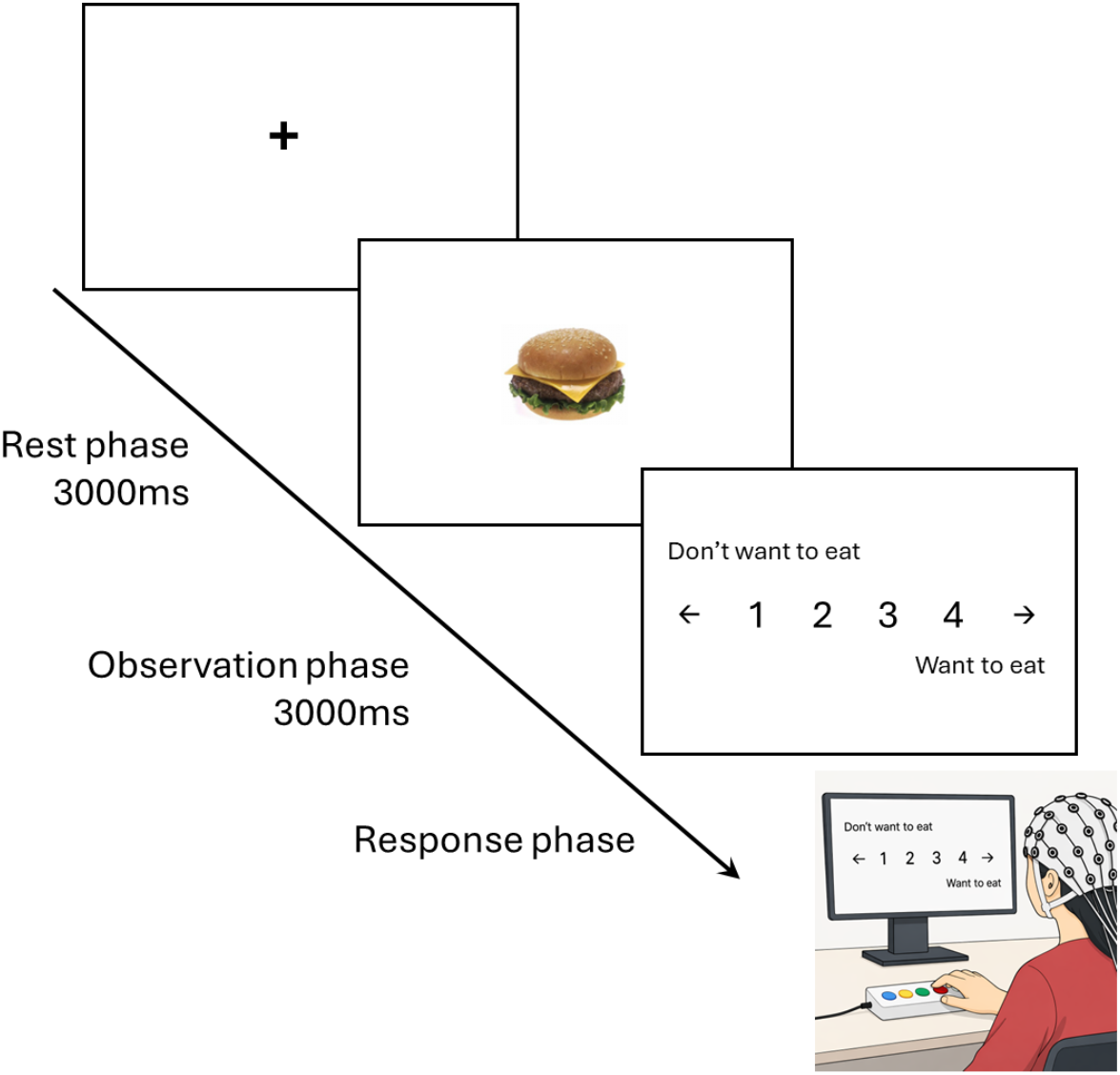
Experimental design. Participants underwent a food observation experiment. Each trial started with a rest phase, during which they were instructed to fixate on a central cross. A food picture was then presented for 3,000 ms, with participants continuously viewing the image. Finally, participants rated their subjective preference for the presented food on a 4-point scale, ranging from 1 (“don’t want to eat”) to 4 (“want to eat”), by pressing one of four buttons.

To reduce EEG artifacts, participants were instructed to minimize unnecessary body movements during the task and maintain visual fixation during the rest and observation phases. For subsequent analyses, trials were classified according to the participants’ responses: ratings of 1 or 2 were categorized as disliked food (DF) trials, whereas ratings of 3 or 4 were categorized as favorite food (FF) trials.

### 2.3 EEG measurements

During the experiment, surface EEG data were recorded to investigate the neurophysiological mechanisms underlying cortical responses to food preferences. EEG data were obtained in an electrically shielded room using a 64-channel EEG system (g. HIamp, g.tec, Austria) at a sampling rate of 1,000 Hz with an online band-pass filter set to 0.1–100 Hz. The ground electrode was placed on the forehead and the reference electrodes attached to the left and right earlobes. Signals from the right hemisphere were referenced to the right earlobe, and those from the left hemisphere referenced to the left earlobe. Active electrodes were positioned according to the international 10–10 system. To monitor eye blinks, electrooculographic (EOG) activity was recorded simultaneously with the EEG. Electrode impedance was maintained <50 kΩ.

### 2.4 EEG data processing

As mentioned above, all trials were divided into two categories for each participant: FF trials and DF trails [6]. Responses of 1 or 2 were assigned to the DF trials, whereas responses of 3 or 4 were assigned to the FF trials. Since the frequencies of the four response options were not identical, the number of trials was balanced before analysis. Specifically, for response pairs 1 versus 4 and 2 versus 3, the smaller trial count was used to match the number of trials within each pair. Then, trials for responses 1 and 2 were merged as DF, and those for responses 3 and 4 as FF. As a result, the final numbers of DF and FF trials were equalized for each participant, and these data were entered into the subsequent analyses.

EEG data were analyzed using Brainstorm software [16, 17]. A notch filter eliminated the power line noise (60 Hz). During the experiment, eye blinking was monitored using an EOG. EEG data were subjected to independent component analysis (ICA; infomax algorithm) to detect regularly occurring artifacts, such as eye blinking and cardiac response. All ICA activations were reviewed on a scrolling display, and components with time courses resembling eye blinks and cardiac artifacts were identified. Second, the nature of these components was validated by plotting their scalp topographies. Identified artifact components were removed from EEG data by back-projecting the remaining components. Then, EEG data were re-referenced to the arithmetic mean of all EEG electrodes (common average).

The recorded EEG signals were time-locked to the onset of food image presentation (0 ms). Noise and data covariances were computed over the 1000-ms periods before and after stimulus onset, respectively. Source reconstruction was then performed using a linearly constrained minimum variance beamformer implemented in Brainstorm with default parameters [18]. In the unconstrained orientation model, each cortical vertex is represented by three orthogonal dipoles aligned along the x-, y-, and z-axes. To derive a scalar time series from these vector-valued source estimates, the Euclidean norm was calculated at each time point as follows:

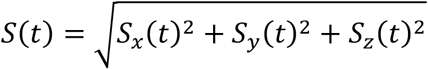

where *S(t)* denotes the scalar magnitude of the source vector at time *t*, computed from the x-, y-, and z-oriented dipole components. This procedure was used to reduce the three-dimensional source estimate at each cortical location to a single-valued time series. Finally, parcel-level signals were obtained by averaging these scalar source values across all vertices within each region of interest defined according to the Desikan–Killiany atlas [19].

### 2.5 Functional connectivity analysis

Functional connectivity (FC) analysis was performed using an ROI-to-ROI approach using Brainstorm with default parameters. To estimate the FC associated with FF and DF trials, amplitude envelope correlation (AEC) was calculated over the 1000-ms period after food-picture onset. Orthogonalization was applied to reduce the pairwise effects of linear signal mixing on long-range temporal correlations and to enable the assessment of relationships between local cortical scaling exponents without artificial coupling between regions [20, 21]. AEC was computed by correlating the amplitude envelopes of two oscillatory brain signals, defined as the absolute value of the Hilbert transform of each cortical oscillation. Higher AEC values indicate more synchronous fluctuations in the amplitude envelopes between oscillations. Thus, AEC can be used to detect synchronization between functional brain networks within specific frequency bands [22-24]. In the present study, the Hilbert transform was applied to band-pass-filtered signals in the alpha (8–13 Hz) and beta (13–25 Hz) bands to obtain complex-valued analytic signals. AEC values were then calculated for each frequency band.

Next, we applied proportional thresholding to create binarized weighted matrices representing reliable connectivity, following established methods (Liu et al., 2014; Sporns et al., 2021). Specifically, we considered the top 10% of FC values between node pairs as reliable connections, as previously reported [25-28]. Finally, we identified significant connectivity links across participants, stronger than the zero-median distribution, using one-sample Wilcoxon signed-rank tests with Bonferroni correction at a significance level of p = 0.05 (Fig. 2) [29, 30].

**Figure 2.**
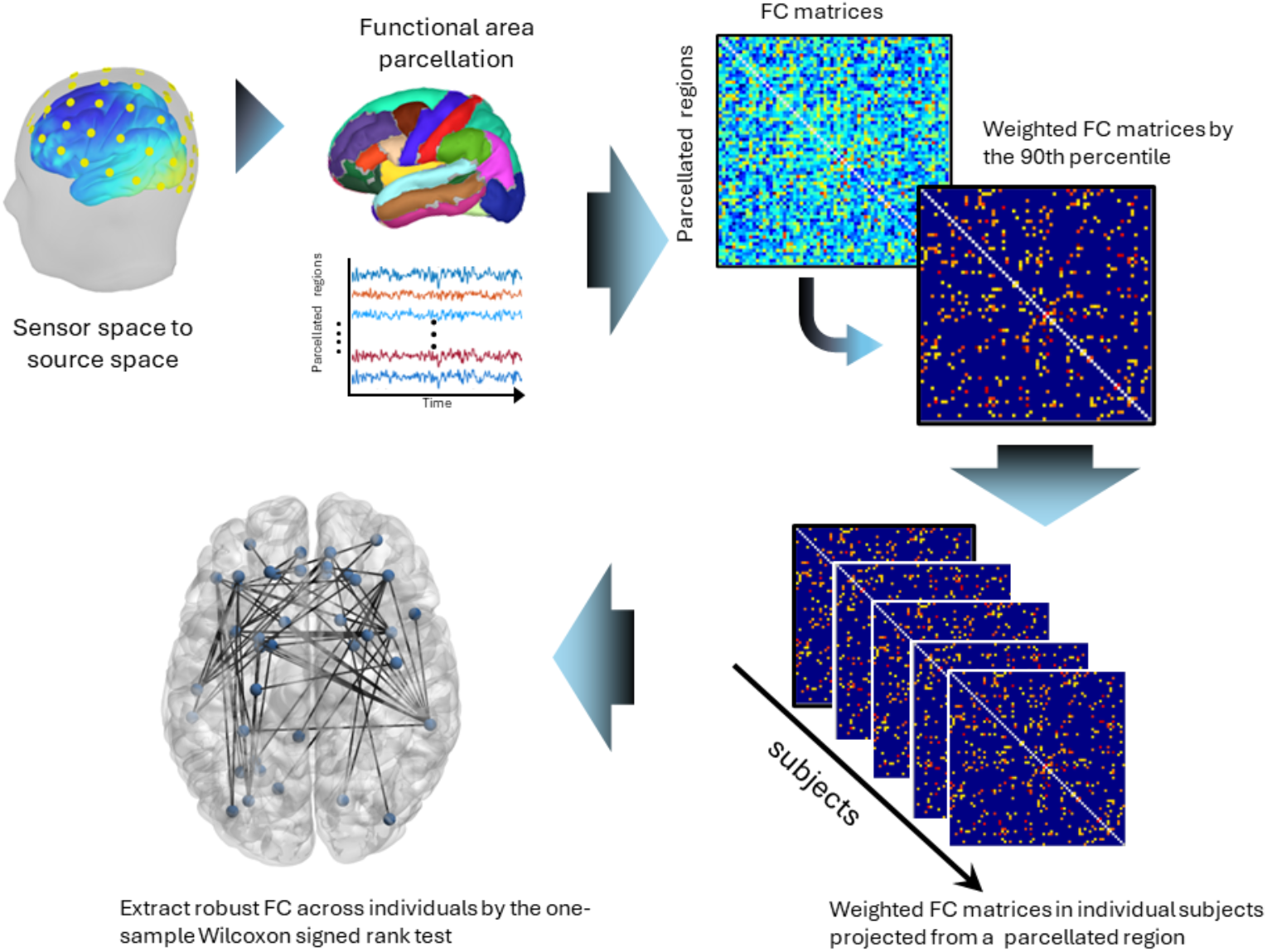
Analysis pipeline for functional connectivity analysis. EEG data were preprocessed and source-localized. Then, source data were parcellated into 60 brain regions based on the Desikan–Killiany Atlas. Functional connectivity (FC) analysis was performed using the amplitude envelope correlation in alpha and beta bands. FC strengths in the top 10% were considered reliable connections. Finally, one-sample Wilcoxon signed-rank tests with Bonferroni correction were performed across participants to identify the most relevant connections related to food preferences.

## 3. Results

### 3.1 Response time

Figure 3 shows the distribution of response times for FF and DF trials. No significant difference in response time was observed between trials, suggesting that the FC was unlikely due to differences in behavioral response speed.

**Figure 3.**
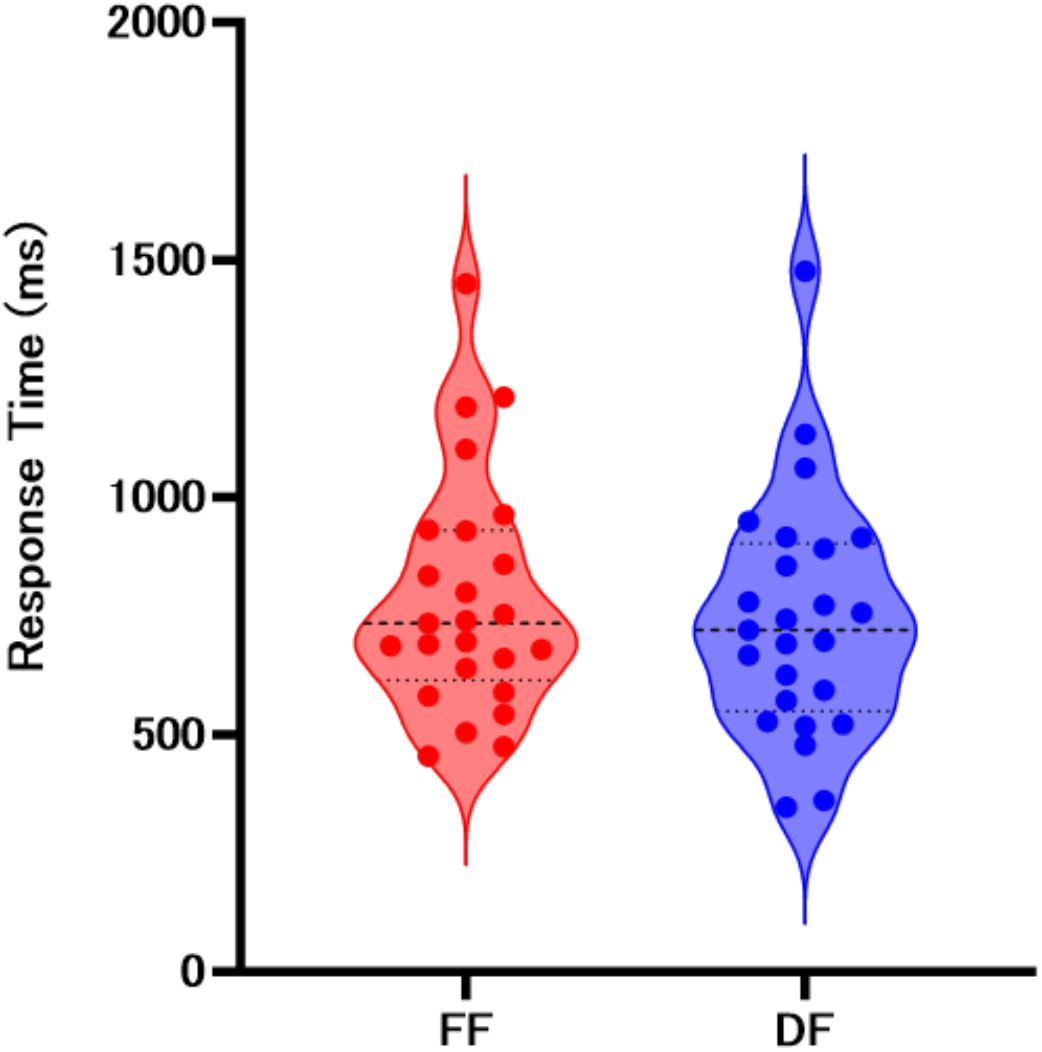
Response times during favorite food (FF) and disliked food (DF) trials. No significant difference in reaction time was observed between the two conditions. Dashed horizontal lines indicate the median and interquartile range.

### 3.2 Functional connectivity in alpha and beta bands

Significant FC was observed in both bands during FF and DF trials (Fig. 4 and 5). In the alpha band, the FF network involved the postcentral gyrus, supramarginal gyrus, cingulate regions, parietal regions, cuneus, and lateral occipital cortex, whereas the DF network involved the isthmus cingulate, caudal middle frontal gyrus, inferior temporal cortex, superior parietal lobule, and lateral occipital cortex (Fig.4).

**Figure 4.**
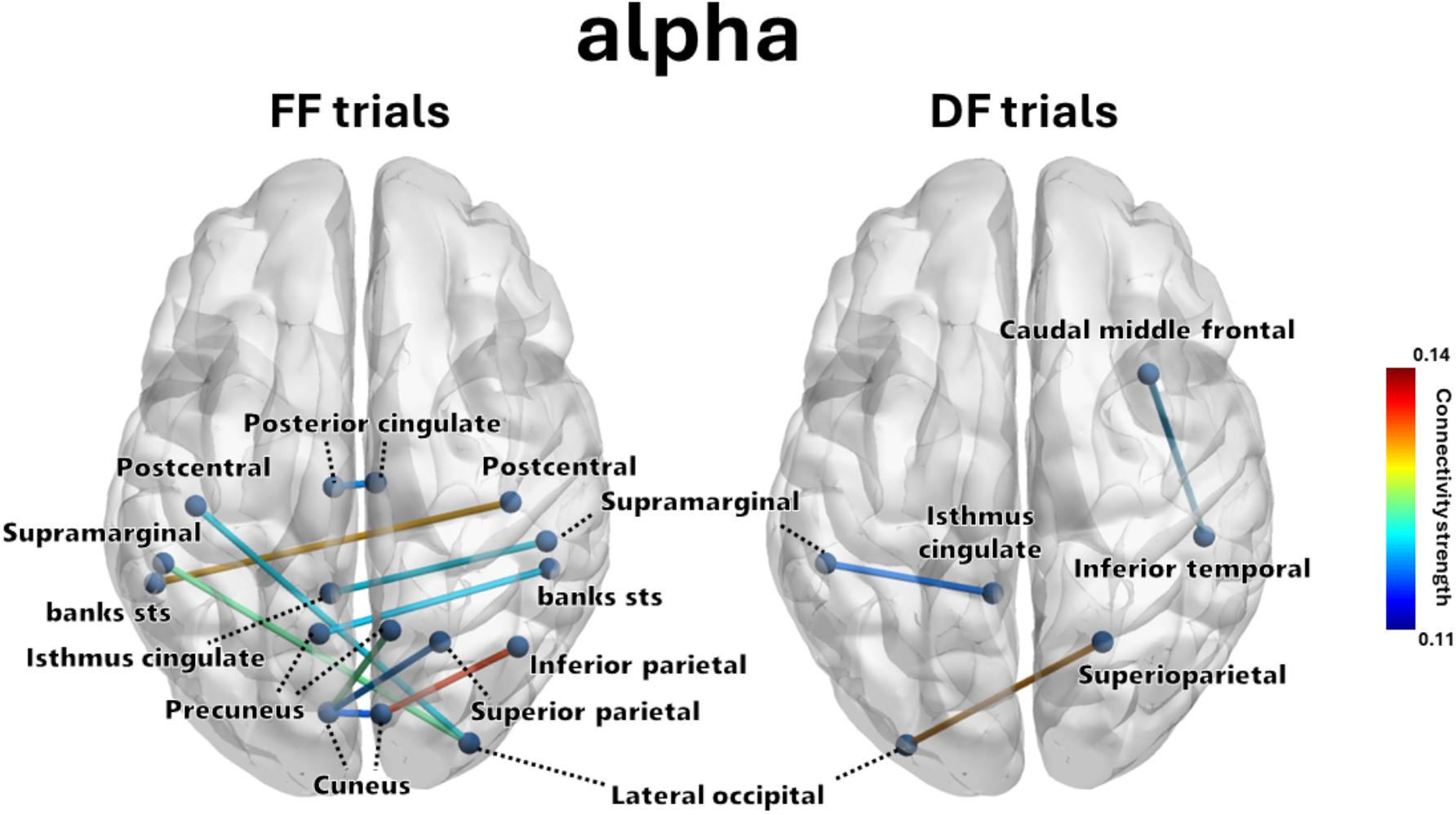
Significant connectivity links in the alpha band during favorite food and disliked food trials. Brain networks showing significant functional connectivity during favorite food (FF) and disliked food (DF) trials in the alpha frequency band. During FF trials, alpha-band connections involved the postcentral gyrus, supramarginal gyrus, cingulate regions, parietal regions, cuneus, and lateral occipital cortex, whereas during DF trials they involved the isthmus cingulate, caudal middle frontal gyrus, inferior temporal cortex, superior parietal lobule, and lateral occipital cortex, with involvement of the caudal middle frontal gyrus in the DF network. Line color indicates average connectivity strength across subjects.

**Figure 5.**
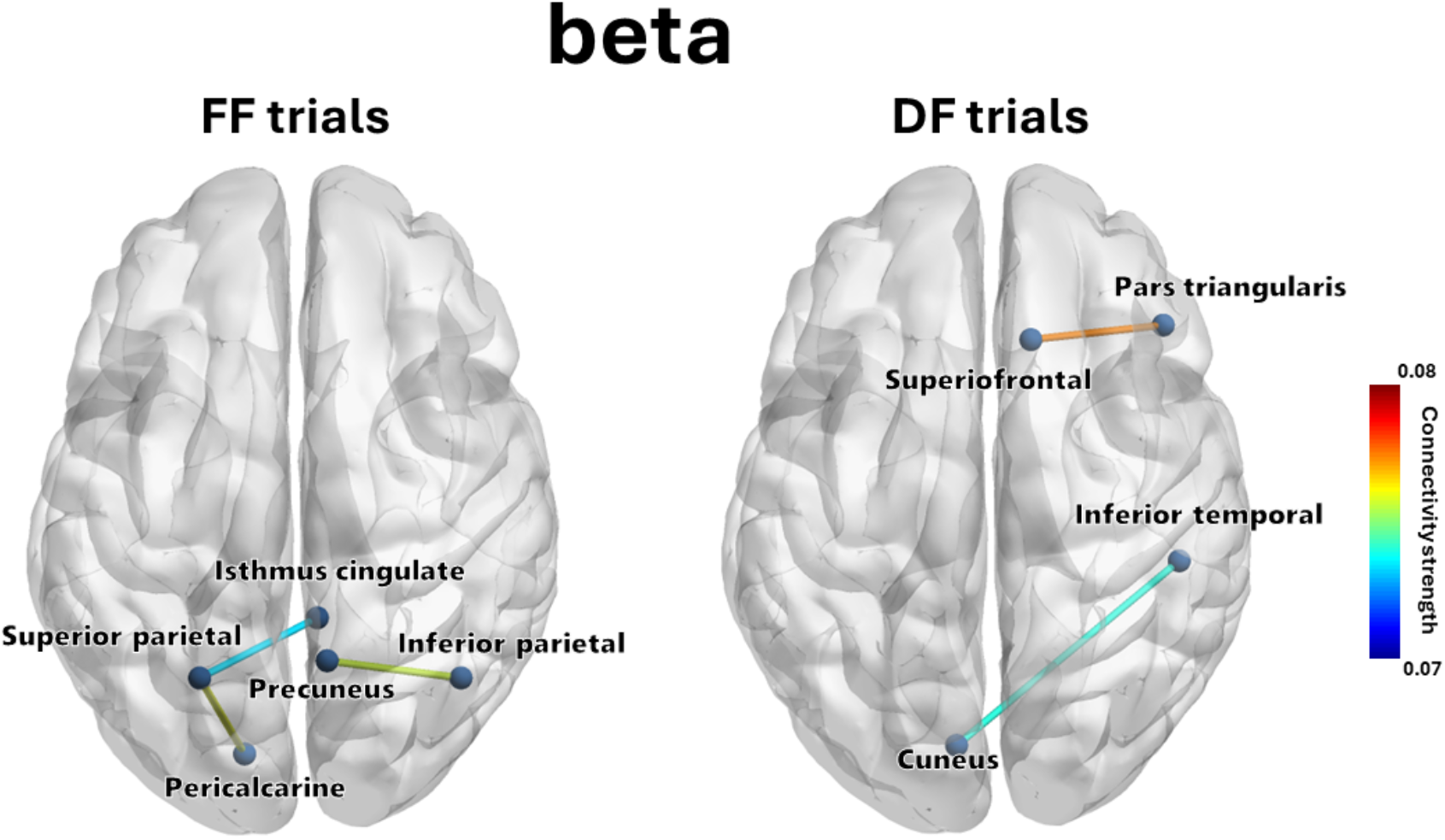
Significant connectivity links in the beta band during favorite food and disliked food trials. Brain networks showing significant functional connectivity during favorite food (FF) and disliked food (DF) trials in the beta frequency band. During FF trials, beta-band connectivity links involved the isthmus cingulate, precuneus, superior parietal lobule, inferior parietal lobule, and pericalcarine cortex, whereas during DF trials they involved the superior frontal gyrus, pars triangularis, inferior temporal cortex, and cuneus, suggesting frontal involvement in the DF network. Line color indicates average connectivity strength across subjects.

In the beta band, the FF network involved the isthmus cingulate, precuneus, superior parietal lobule, inferior parietal lobule, and pericalcarine cortex, whereas the DF network involved frontal regions, including the superior frontal gyrus and pars triangularis, as well as the inferior temporal cortex and cuneus (Fig. 5). These results indicate that FC patterns differed between FF and DF trials in both the alpha and beta frequency bands, with the DF condition showing more prominent frontal involvement.

## 4. Discussion

The present study revealed that FF and DF trials were associated with distinct FC patterns during food image observation. Although response time did not differ between conditions, the alpha- and beta-band networks differed between FF and DF trials. These findings suggest that food preference is represented at the level of large-scale cortical network organization during food-cue evaluation.

A main finding of the present study was the prominence of alpha-band connectivity during FF trials, particularly in networks involving the cuneus, lateral occipital cortex, parietal regions, and cingulate regions. This pattern is consistent with previous studies showing that alpha-band activity is closely related to sensory processing, perceptual gating, and attentional allocation during early stimulus evaluation [9, 10]. Visual food cues engage cortical systems involved in attention, evaluation, and motivational processing before actual intake occurs([3, 5, 31, 32]. In this context, the present FF alpha-band network may reflect a cuneus-centered visuo-attentional system involved in prioritizing subjectively desirable food cues. The involvement of parietal regions may reflect attentional orienting and multimodal integration, while cingulate involvement may be related to the evaluative significance of the stimuli [32-35]. Thus, the FF alpha-band network may capture an early stage of food-cue evaluation in which desirable food images are preferentially processed through distributed visual-attentional mechanisms. In addition, the beta-band connectivity observed during FF trials involved the isthmus cingulate, precuneus, parietal regions, and pericalcarine cortex, suggesting integration of visual information with internally oriented and sensorimotor-related representations, as beta oscillations have often been associated with sensorimotor and evaluative processing [7, 12, 13, 28].

This interpretation extends previous EEG findings showing that food preference modulates oscillatory brain activity [6]. Earlier work reported localized oscillatory changes, whereas the present study suggests that food preference is also expressed through distributed cortical interactions. By using high-density EEG and source-level ROI-to-ROI FC analysis, the present study expands the neurophysiological account of food preference from local activity changes to large-scale networks. The prominence of alpha-band effects suggests that food preference may be expressed during passive viewing of food images, particularly through sensory-attentional prioritization rather than through localized activity changes alone [9, 10]. This interpretation is consistent with the view that low-frequency oscillations support large-scale cortical communication and gating during perceptual and cognitive processing [36, 37].

In contrast, the DF condition showed a different pattern. In the alpha band, connectivity involved the isthmus cingulate, caudal middle frontal gyrus, inferior temporal cortex, superior parietal lobule, and lateral occipital cortex, while in the beta band, the network also involved frontal regions, including the superior frontal gyrus and pars triangularis. This pattern suggests that DF cues engage a more selective, evaluative mode of processing than preferred food cues. Rather than reflecting approach-related attentional engagement, nonpreferred food images may require additional discrimination or rejection-related processing. Food cues do not simply vary in hedonic value but also in the extent to which they elicit motivational salience and prompt approach or avoidance tendencies [38]. In the present context, the frontal involvement observed in the DF network is compatible with higher-order evaluative control during the judgment that a food is not desirable. Because our task evaluated subjective preference rather than explicit disgust, these findings are better interpreted as reflecting nonpreferred or rejection-related processing rather than disgust itself.

Taken together, the alpha- and beta-band findings suggest that food preference influences multiple stages of food-cue evaluation, with partially distinct roles across frequency bands. Alpha-band networks may reflect early visual-attentional prioritization of food cues, whereas beta-band networks may reflect subsequent integration with internally oriented, sensorimotor-related, and evaluative processes. Within this framework, preferred food cues may enhance attentional engagement, whereas disliked food cues may involve greater frontal evaluative processing. These findings suggest that food preference shapes both the early salience of food cues and their later appraisal in relation to desirability.

From the perspective of food choice research, these findings indicate that subjective preference is associated with distinct patterns of cortical connectivity even before overt eating. This may help understand how the brain organizes food-related decisions before actual selection or intake [2, 3, 5, 32]. FC analysis may therefore provide a useful framework for investigating individual differences in food preference and cue-driven eating-related behavior.

Several limitations should be acknowledged. First, the sample size was modest after excluding participants with highly unbalanced response patterns, which may limit the generalizability of the findings. Second, the classification of trials into FF and DF was based on subjective ratings, and the DF condition likely included heterogeneous states such as low appetite, indifference, or aversion. Third, although source-level EEG connectivity analysis offers high temporal resolution, its spatial resolution remains limited. Fourth, because the task involved only visual food-cue observation, the findings cannot be directly generalized to actual food choice behavior or intake. Future studies should examine whether similar connectivity patterns predict real-world eating decisions, food selection, or intake-related behavior. It will also be important to determine whether these network patterns vary according to hunger state, caloric value, or other motivational features of food cues.

In conclusion, the present study demonstrated that food preference is associated with distinct large-scale cortical FC patterns during food image observation. Preferred food cues were characterized by a cuneus-centered alpha-band visuo-attentional network involving parietal, cingulate, and occipital regions, whereas disliked food cues showed relatively greater frontal involvement, particularly in the beta band. These findings suggest that food preference influences early stages of food-cue evaluation through distributed cortical networks related to visual-attentional and evaluative processing.

## CrediT Authorship contribution statement

**Hisato Sugata:** Writing – review & editing, Writing – original draft, Supervision, Project administration, Methodology, Investigation, Formal analysis, Data curation, Conceptualization. **Sarang Kim:** Data curation. **Takashi Ikeda:** Software. **Masayuki Hrata:** Methodology and Conceptualization.

## Ethical statement

All procedures were performed following the Declaration of Helsinki. Prior to participation, all participants received a full explanation of the purpose of the study and its potential consequences, and written informed consent was obtained from each participant. The study protocol was reviewed and approved by the Ethical Review Board of the Faculty of Welfare and Health Science, Oita University.approval no. F240009).

## Data availability

Behavioral and EEG data are available upon request by contacting the corresponding author, Hisato Sugata.

